# MEGA-FISH: multi-omics extensible GPU-accelerated FISH processing framework for huge-scale spatial omics

**DOI:** 10.1101/2024.12.04.626913

**Authors:** Yuma Ito, Kosuke Tomimatsu, Masao Nagasaki, Hiroshi Ochiai, Yasuyuki Ohkawa

## Abstract

Spatial omics enables comprehensive mapping of cell types and states in their spatial context, providing profound insights into cellular communication and tissue organization. However, analyzing large tissue sections, especially crucial for clinical applications, remains a significant challenge due to the computational demands of current image processing methods. To overcome these limitations, we developed MEGA-FISH, a flexible, GPU-accelerated Python framework optimized for large-scale spatial omics image analysis. Benchmarking on simulated and tissue images demonstrated that MEGA-FISH achieved high accuracy in spot detection while significantly reducing processing times compared with established tools. The framework’s adaptable computational capabilities optimize resource allocation (e.g., GPU or multi-core CPU) for diverse tasks, and its scalable architecture enables integration with advanced imaging and segmentation techniques. By bridging cutting-edge imaging methods and single-cell analysis, MEGA-FISH provides an efficient platform for multi-modal analysis and advances research and clinical applications of spatial omics at organ and organism scales.

## INTRODUCTION

In pathological diagnosis, cell states are evaluated based on the expression levels of biomolecules, such as RNA and proteins, to identify lesion sites within large histological sections. Recent advancements in spatial omics technologies have enabled the comprehensive analysis of cell states associated with disease pathogenesis^1,2^. Imaging-based techniques, such as single-molecule RNA fluorescence in situ hybridization^3–5^ and immunofluorescence^6^, have been developed to capture the abundance and spatial distribution of these biomolecules in situ. These methods have been enhanced by sequential imaging^7,8^, enabling the determination of cell types from their expression patterns. Recent multiplexed approaches^9,10^, combined with advanced high-dimensional image processing^11,12^, enable comprehensive lineage tracking and cell fate mapping^13,14^. These advancements have propelled the integration and expansion of spatial multi-omics analyses at organ- and organism-wide scales. At these scales, a single experiment can generate terabytes of image data, with computational costs hundreds of thousands of times higher than methods that target a small number of cells. These challenges underscore the need for transformative high-throughput processing of huge-scale images.

Imaging-based spatial omics uses various specialized processing steps, including registration, cell segmentation, and spot detection. Established methods, such as Big-FISH^15^ and RS-FISH^16^, have simplified complex spot detection tasks and achieved nanometer-scale localization precision. However, these methods cannot process images that exceed memory capacity and require external batch processing setups to support parallel computing. Barcoding-based methods, such as MERFISH^11^, seqFISH+^10^, and starfish^17^, have the potential to leverage Graphics Processing Unit (GPU) acceleration, driven by recent advancements in hardware performance. To perform simultaneous analysis of spot detection and methods such as immunofluorescence, pipelines for spatial multi-omics analysis are available^18,19^. However, these pipelines are often tightly coupled with specific experimental protocols, posing challenges for integration with emerging techniques (summarized in Supplementary Table 1). Therefore, a framework that allows flexible selection of both computational resources and imaging modalities is essential for advancing spatial multi-omics analysis.

To meet this demand, we developed the MEGA-FISH (Multi-omics Extensible GPU-Accelerated-Fluorescence In Situ Hybridization) processing framework, a flexible Python-based image processing package. This framework allows users to select and allocate computational resources, such as GPUs and CPUs, for specific processing tasks. With its step-by-step workflow that uses a simplified data structure, MEGA-FISH facilitates the efficient processing of large-scale images, reducing the complexity of handling deeply multiplexed and multi-modal images. MEGA-FISH enables tasks such as incorporating new imaging techniques and data preparation for state-of-the-art external segmentation tools.

## RESULTS

### Scalable Framework for Large-Scale Spatial Multi-Omics Analysis

We developed a Python-based image analysis framework, MEGA-FISH, for efficient processing of large-scale spatial multi-omics data (Fig. 1). The workflow begins by registering tile-based images captured by seqFISH or sequential immunostaining (SeqIS) into a single large-scale xarray data structure^20^. This approach eliminates the need for users to manually organize file names and define field-of-view relationships. Subsequent modality-specific processing, including spot detection and protein intensity calculation, is performed on each chunk of the data structure using Dask for parallel computing, and CuPy and RAPIDS are leveraged for GPU computation. These resources can be switched dynamically depending on the task, enhancing processing speed without additional Spark-based wrapping or batch processing. These approaches collectively streamlined the workflow, lowering the user entry barrier while maintaining flexibility in data management. The modular structure of MEGA-FISH also facilitates high extensibility, enabling fine-tuning for specific modalities and individual observations. We ensured high accessibility and flexibility by storing intermediate data in Zarr format and using Napari^21^ for visualization, thus allowing for simultaneous data validation and adjustment. These functionalities ensure that MEGA-FISH provides a scalable, accessible, and adaptable framework for processing large-scale spatial multi-omics images.

**Figure 1.**
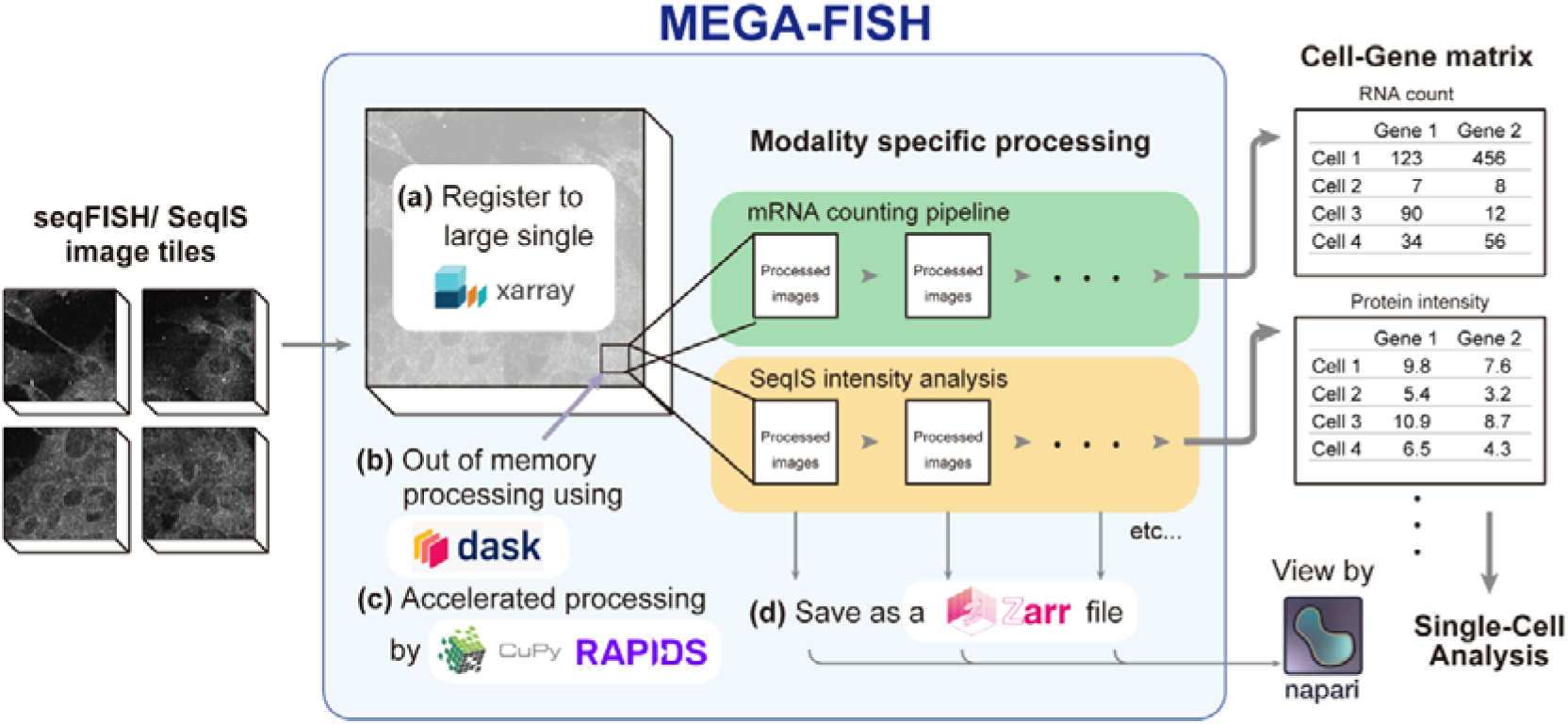
Overview of the data processing workflow in MEGA-FISH. Sequential and tiled images are assembled into a large xarray data structure (a), and each chunk is processed in parallel using Dask (b). Modality-specific processing is run in submodules with GPU acceleration (c). All processed images are saved in Zarr format and visualized with Napari (d), and the resulting cell–gene matrix is used for single-cell analysis.

### Spot Detection Reproducibility and Performance

To evaluate the reproducibility of MEGA-FISH and established analysis programs, we calculated spot detection precision using simulated images containing randomly distributed theoretical fluorescence spots (Supplementary Fig. 1a). We compared the calculation speed from image loading to the generation of spot coordinates for MEGA-FISH, Big-FISH, and RS-FISH, which we executed with a minimum configuration, without additional parallelization using Spark or batch processing. MEGA-FISH processed the images 3.5 times faster than Big-FISH pixel-level detection (Fig. 2a). To quantify the spot detection performance, we calculated the localization error and Jaccard index, which measure positional accuracy and detection completeness, respectively (Fig. 2b). The root mean square error between detected spots and the ground truth positions across varying densities showed sub-diffraction accuracy in MEGA-FISH (Fig. 2c), consistent with Big-FISH pixel-level detection, suggesting sufficient precision for spot counting per cell. The detection performance of MEGA-FISH was equivalent to those of Big-FISH and RS-FISH at a wide range of spot densities, achieving high accuracy at densities up to approximately one spot/µm² (Fig. 2d). To confirm reproducibility in tissue samples, we analyzed previously published human uterine carcinosarcoma data^22^. The sections were processed into a single stitched image before analysis (Supplementary Fig. 1b). RNA counts showed strong correlation across different tissue sections (Fig. 2e). Comparison of mean RNA counts per cell showed strong correlations between MEGA-FISH and both Big-FISH and RS-FISH (Fig. 2f, g). These results suggest that MEGA-FISH provides robust and consistent detection across varying conditions of biological samples.

**Figure 2.**
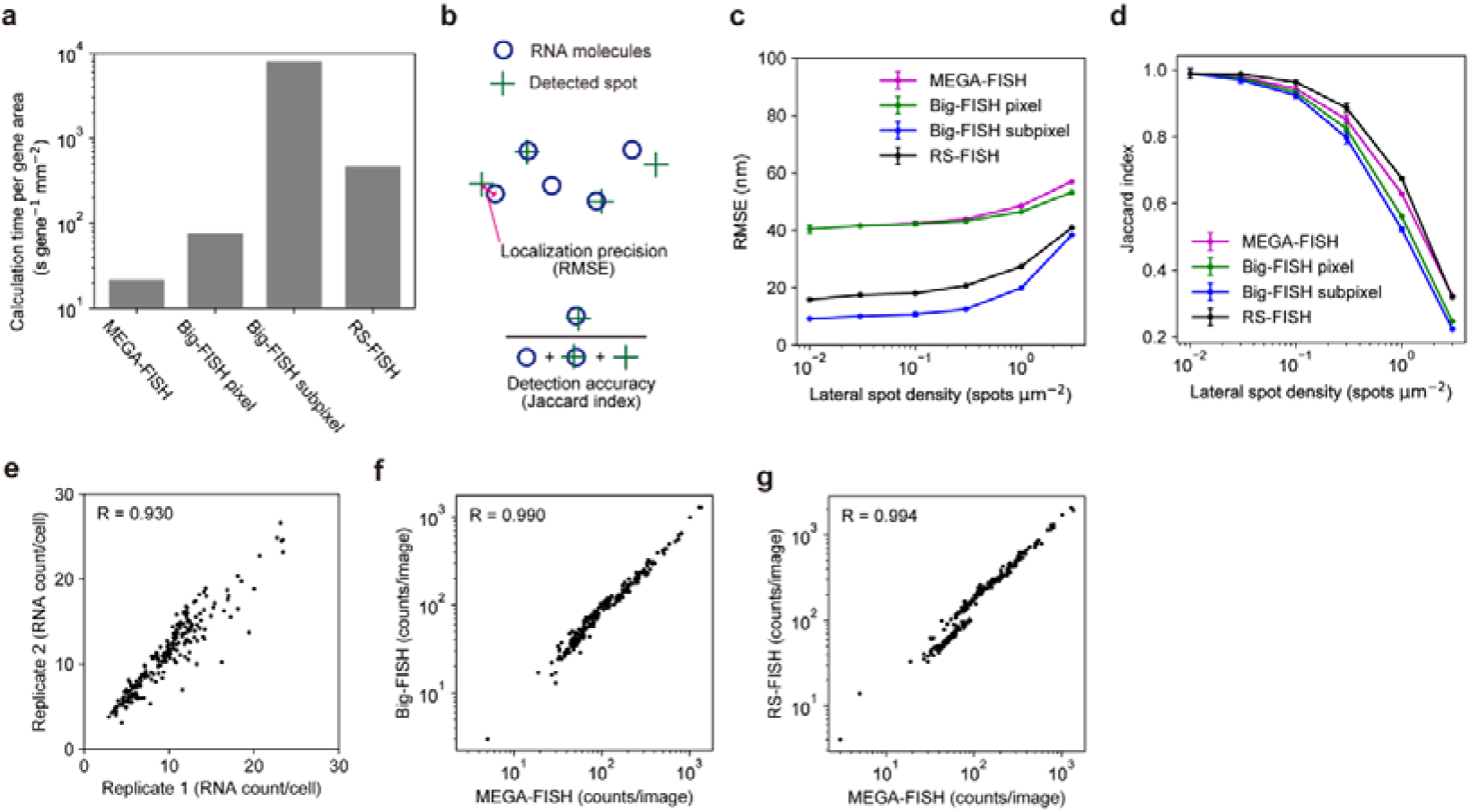
Evaluation of spot detection accuracy and reproducibility for RNA-seqFISH. (a) Normalized computation time for detecting simulated spots using different methods. (b) Schematic illustration of localization precision and detection completeness. (c) Root mean square error (RMSE) between the ground truth positions and true positive detections. Spots within 100 nm of the ground truth are considered true positives. (d) Detection accuracy of simulated spots across varying spot densities. (e) Reproducibility of spot counts per gene between two tissue sections. Each dot represents the average spot count for each of the 220 genes. (f, g) Comparison of RNA-seqFISH spot counts per gene in a single field of view compared with the counts for Big-FISH (f) and RS-FISH (g).

### Factors that Influence Computational Efficiency in MEGA-FISH

Given that spot detection efficiency of MEGA-FISH was compatible with established methods, we investigated the computational configuration that affected the efficiency of the entire process, from loading images to spot counting (Fig. 3a). We used a desktop computer equipped with a multi-core CPU and a non-enterprise GPU to perform benchmarking. We assessed the image loading speed by comparing different CPU configurations (synchronous, multi-thread, and multi-process) with varying storage types (hard disk drive, solid state drive, M.2 solid state drive) (Fig. 3b). We normalized the computation time to the time required to process one mm² area per gene. The normalized calculation time showed that the multi-process CPU setup had the fastest loading times and that the differences between storage types were minimal, indicating that the primary bottleneck was likely the handling of compressed images. Next, we evaluated the efficiency of registration processing using phase cross correlation, scale invariant feature transform (SIFT)^23^ and stitching across different computational units (Fig. 3c). The GPU had the fastest processing speed for phase cross correlation, but the advantage of the GPU was less pronounced for SIFT and stitching, indicating that the effectiveness of GPU acceleration may be task dependent. For seqFISH-specific spot detection, the optimized counting algorithm of MEGA-FISH provided faster processing than other processing steps, including loading and registration (Fig. 3d), suggesting a minor effect on the whole processing time. The performance gains from multi-process and GPU configurations were limited, likely because of the increasing overhead for parallel computing. We also investigated the decoding process of the barcoded multiplex seqFISH method using simulated codebooks (Fig. 3e). As codebook size increased, the processing time grew linearly for CPU-based operations, resulting in exponential growth with the number of imaging cycles. The GPU provided high-speed processing and achieved a speed increase of up to 1000-fold compared with that of a single CPU. We examined the impact of chunk size during pixel decoding (Fig. 3f). Regardless of the configuration, small chunk sizes resulted in increased processing time. Although large chunk sizes improved speed, available random-access memory (RAM) size constraints limited the feasibility of this approach. These results highlight the importance of selecting the appropriate computational configuration to optimize processing speed. The flexible resource allocation of MEGA-FISH enables users to choose the optimal configuration for each task.

**Figure 3.**
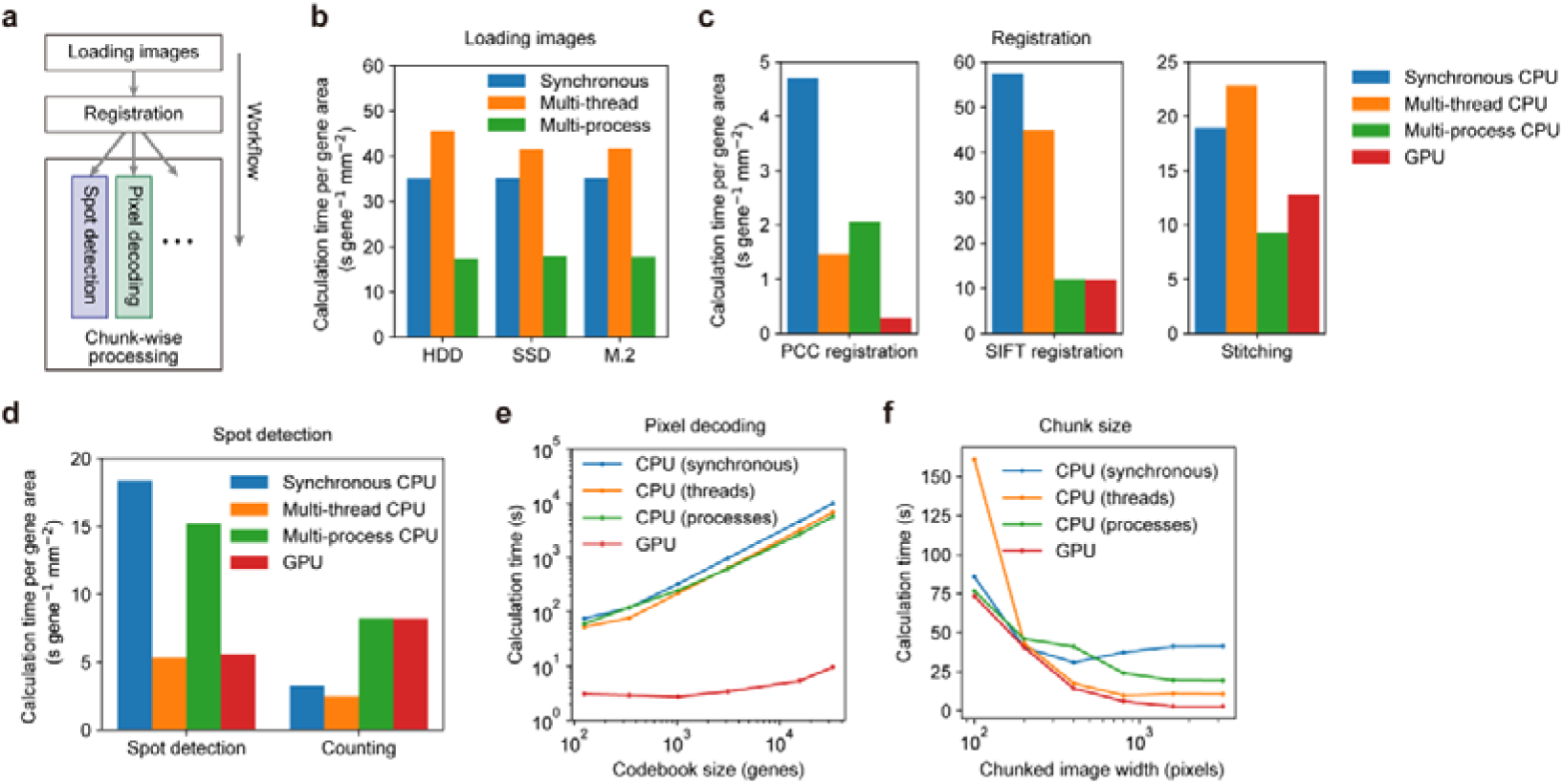
Evaluation of computational performance across various settings for image processing. (a) Schematic illustration of the seqFISH processing steps. (b) Computation time for loading images in Zarr format across different CPU and storage configurations. Processing time was normalized to the number of imaging channels, area, and cycles. (c) Computation time for image registration using different CPUs and a GPU. The CPU scheduler selected the fastest processing method for each task to optimize GPU-based calculations. (d) Computation time for seqFISH-specific processing steps. (e) Effect of codebook size on processing speed during nearest neighbor decoding for pixel-level gene determination. (f) Effect of chunk size on processing performance during pixel vector decoding. The decoding step includes pre-filtering and nearest neighbor decoding.

### Extension of MEGA-FISH with SeqIS and Segmentation Methods

Spatial omics technologies are continuously being developed; thus, the framework should have scalability to adapt to novel imaging and analysis methods. To demonstrate the flexibility of the MEGA-FISH pipeline, we incorporated the latest SeqIS method and segmentation tool into the workflow (Fig. 4a). For this, we created a SeqIS-specific module suitable for PECAb^22^, an antibody linked to a fluorescent dye via a disulfide bond (Fig. 4b). The immunofluorescence signal is precisely erased using the reducing reagent TCEP [tris(2-carboxyethyl)phosphine], and subtracting TCEP-treated background images to further enhances protein signals. We also added scripts for fine-tuning input images for MEDIAR^24^ segmentation (Fig. 4b), in which we selected a combination of cell membrane markers, supplemented by α-smooth muscle actin (αSMA) for cells with low marker expression (Fig. 4c)^25^. The segmented labels showed increased performance of cell detection and a broader segmentation area per cell compared with nuclear-only segmentation with Cellpose (Fig. 4d). The post-segmentation process enabled seamless integration of labels across chunks (Fig. 4e) and yielded a 1.4-fold increase in cell detection (Fig. 4f). The normalized expression distribution of representative proteins showed that the analysis using TCEP image subtraction combined with MEDIAR highlighted the differences in expression levels between cells more efficiently than the analysis using Cellpose segmentation of fluorescence images alone (Fig. 4g). The distribution of cell features after dimensionality reduction by principal component analysis showed a more dispersed distribution (Fig. 4h) in the major population. The standard deviation of principal components showed a slight increase in variance along the first component with significant reductions in variance across subsequent components (Fig. 4i). This separation of principal components aligns with the presence of two major cell types, epithelial and mesenchymal cells, in the tissue sections. These results suggest that because of the flexibility of MEGA-FISH, state-of-the-art techniques can be integrated into the workflow and can improve the separation of cell features in spatial single-cell analysis.

**Figure 4.**
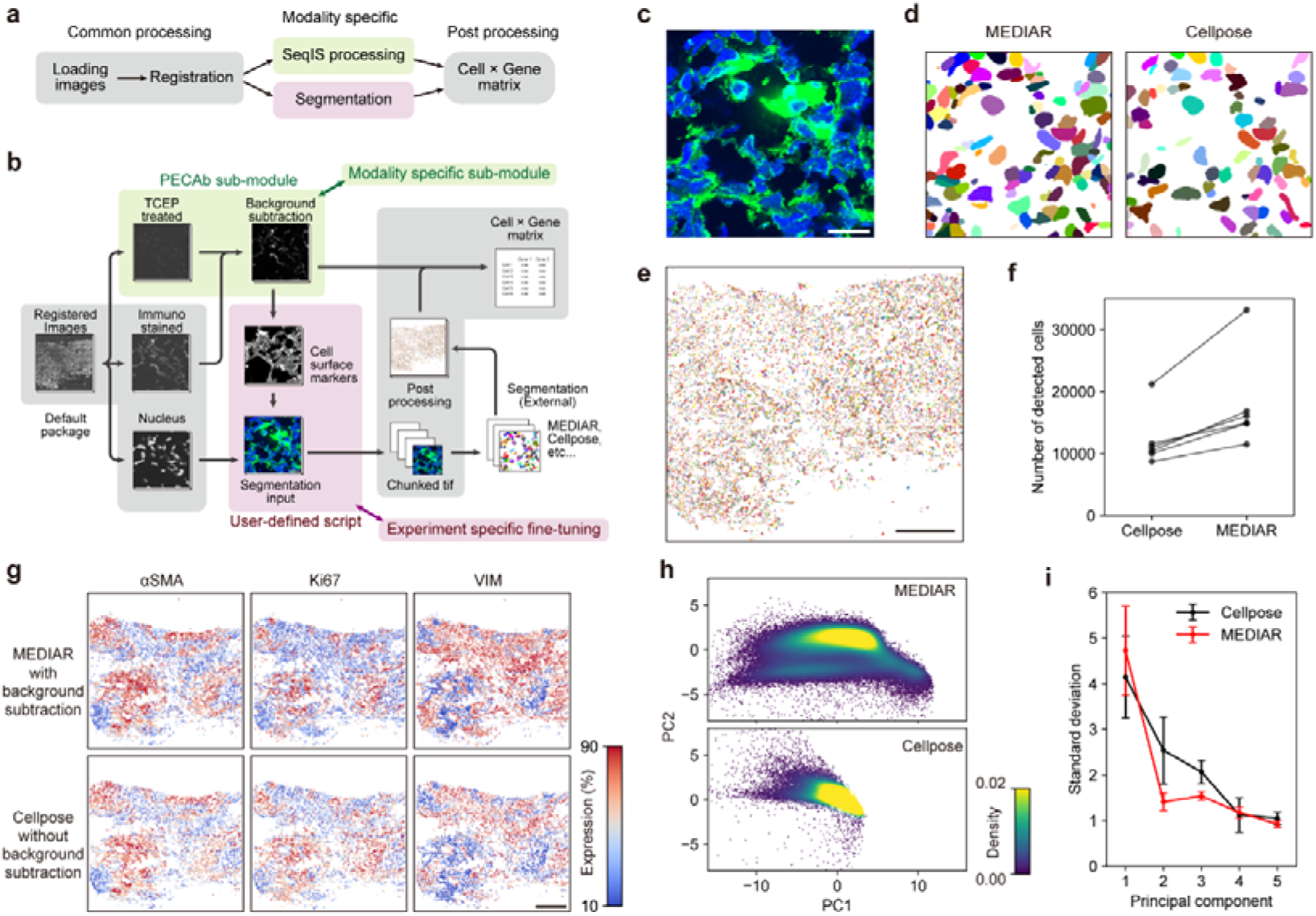
Extended capabilities of the MEGA-FISH pipeline with sequential immunostaining (SeqIS) and segmentation methods. Overview (a) and detailed (b) schematic of SeqIS analysis workflow using the PECAb-specific module and user-defined scripts for segmentation. (c) Representative image showing a chunk of the input used for MEDIAR segmentation. Nuclei are stained with Hoechst 33342 (blue), and the cytoplasm is labeled with protein signals for E-cadherin, N-cadherin, β-catenin, and α-smooth muscle actin (all shown in green). (d) Cell areas segmented using image (c) as input. MEDIAR segmentation was performed on nuclear and cytosolic channels (left). Cellpose segmentation is applied to the nuclear channel (right). (e) Whole tissue section segmentation using MEDIAR. (f) Number of cells detected per tissue section by Cellpose and MEDIAR. (g) Spatial distribution of normalized fluorescence intensities for representative proteins across the tissue sections. MEDIAR segmentation with background subtraction using TCEP-treated images (top row). Cellpose masks were applied to the original fluorescence images (bottom row). Each dot represents the z-scored fluorescence intensity per cell. αSMA, α-smooth muscle actin; Ki67, marker of proliferation Ki-67; VIM, vimentin. (h) Principal component analysis of the expression levels of 35 proteins. The scatter plot shows individual cells in the space of the first two principal components (PC1 and PC2), focused on the major population. (i) Standard deviation of the first five principal components for each tissue section. Error bars represent the standard deviation across six tissue sections. Scale bars: (c) 20 µm, (e) 400 µm, (g) 400 µm.

## DISCUSSION

We developed MEGA-FISH, a Python framework designed to efficiently analyze large-scale spatial omics images by leveraging GPU computing and a scalable architecture. MEGA-FISH achieved significantly accelerated processing speed and maintained an image-processing performance comparable to those of existing tools. The flexible architecture of MEGA-FISH enables the integration of customizable modules for advanced SeqIS methods and segmentation, improving the resolution of cell states within tissue samples.

The ability to adjust computational resources in MEGA-FISH enables the identification of optimal configurations for various processing steps. While GPU acceleration dramatically enhanced decoding and image correlation tasks, specific processes required CPU computation. The CPU-dependent tasks still benefited from parallel CPU calculation, which allowed MEGA-FISH to perform effectively in environments without GPU support or with CPU-based high-performance computing clusters. MEGA-FISH also supports the widely used GPUs, ensuring compatibility with standard computational setups without the need for expensive enterprise-level systems. Point-based registration^26^ and handling of compressed images^27,28^ are processes where GPU support is anticipated. Although the current version of MEGA-FISH is limited to two-dimensional images, expanding to three-dimensions is expected to maximize GPU efficiency due to larger data sizes. The ongoing updates to MEGA-FISH are supported by community-driven development on GitHub where contributions from the community are encouraged.

As demonstrated by the improvements of cell feature identification achieved by combining the PECAb method and MEDIAR segmentation, optimizing the sample staining and image acquisition stages enhanced the spatial omics performance; however, these optimizations are challenging to implement in standardized commercial omics platforms^29–31^. Such modifications are crucial for advancements in imaging-based spatial omics, where novel techniques are continuously being developed in both imaging^19,32^ and analysis^33,34^. With its high flexibility, MEGA-FISH has the potential to become a standard framework, bridging gaps between fluorescence images captured with emerging methods and subsequent spatial^35–37^ and single-cell analysis^38,39^.

The chunk-based analysis of reconstructed multi-modal images not only eliminates the need for users to manage tile positioning and distribution among processing units but also allows for optimization based on state-of-the-art computational environments, such as high-performance GPU and large RAM and storage. The scalability of MEGA-FISH enables the future integration of highly multiplexed spatial omics and whole slide imaging^40^. These advancements facilitate comprehensive analysis of cellular communication at the tissue and organism levels, enabling precise cell state determination and supporting high-precision pathological diagnostics.

## METHODS

### Fluorescence spot simulation

The three-dimensional confocal model was generated using the Python psf package with pixel size 10 nm, numerical aperture (NA) 1.4, and wavelength 519 nm. To simulate realistic imaging conditions, point spread function images were generated from randomly distributed coordinates spaced 10 nm apart, then resized to 100 nm per pixel and embedded into 30 z stacks of 1000×1000 images. Gaussian noise and spot intensities were added based on parameters derived from images captured using a Zyla-4.2 Plus camera (ANDOR Technology, UK).

### Image datasets

The previously published image dataset of human carcinosarcoma tissue^22^ was used for spot detection validation. A 1000×1000 pixel region was extracted from the seqFISH tissue images across all cycles and used to compare the software. Pixel vector decoding was applied to 16-cycle cell images from the starfish tutorial^17^, which were cropped to 1600×1600 pixels and tiled to form a 3200×3200 pixel image. Nearest neighbor calculations for pixel vector decoding were performed using uniform random images with values of 0–1 across all rounds and cycles. The resulting 2000×2000 two-dimensional images were then split into four 1000×1000 pixel chunks for further analysis.

### Image processing

Field of view images were loaded as Zarr files compatible with xarray, and organized into six dimensions (cycle, tile y position, tile x position, z, y, x) for each channel. A z-maximum intensity projection was applied to each tile. Lateral shifts between cycles were initially calculated from Hoechst 33342 images using the phase_cross_correlation function from the scikit-image package^41^. The shifts were further refined by point pair matching based on SIFT, match_descriptors, and RANSAC functions of scikit-image. The position relative to the entire image was calculated using phase cross correlation and SIFT against a pre-stitched reference image. The reference image was manually stitched using external tools Imaris Stitcher (Oxford Instruments, UK) and Image Stitching plugin in Fiji^42^. Homography transformation coefficients were generated for both tile positions and cycle shifts. Tiles in each chunk were merged using maximum intensity projection, forming a large chunked xarray array.

Nuclear and cytoplasmic segmentation was performed on the RGB TIFF (Tagged Image File Format) images for each chunk using the fine-tuned MEDIAR models^24^. For comparison, nuclear segmentation was also performed separately using the nuclear model from Cellpose^43^. Segmentation labels were saved in Zarr format and post-processed to ensure globally unique cell IDs, merging IDs across chunks and filling holes to eliminate gaps.

For spot detection, a Difference of Gaussian filter was applied to the maximum intensity projection images to enhance RNA spots. Spots were identified by calculating local maxima, and the resulting sparse intensity images contained zeros except at local maxima. An intensity threshold was applied to select final spots, which were then assigned to segmented cells by referencing the segmentation label images. For reproducibility, all analysis parameters are available from the publicly accessible GitHub repository.

### Benchmarking

Benchmark analyses were conducted on a desktop workstation equipped with an Intel Xeon E5-2680 v3 CPU (12 cores, 24 threads), 64 GB RAM, and a GeForce RTX 4090 GPU, running the Ubuntu 22.04.5 LTS operating system. Big-FISH version 0.6.2 was used with Python 3.6, and RS-FISH was used as a Fiji plugin in an ImageJ macro. For each tool, maximum intensity projection TIFF images generated by MEGA-FISH were used. Key parameters, including the intensity threshold, were set manually. Spot detection accuracy was computed using the Jaccard index function^44^. Computation times were measured from the point of TIFF image loading to the output of spot coordinates. CPU resource management in MEGA-FISH was handled using Dask, switching the concurrent.futures scheduler provided by Python. For GPU-based computations, the most efficient CPU scheduler was employed. Calculation speed measurements were performed multiple times, but the variability between runs was negligible, so representative times are shown (Fig. 3). Large-scale analysis was performed using the Omics Science Center Secure Information Analysis System (Medical Institute of Bioregulation, Kyushu University).

## Data availability

Custom scripts used in this study are available on GitHub at https://github.com/yumaitou/megafish-scripts. Example datasets for use with MEGA-FISH are available on Zenodo at https://doi.org/10.5281/zenodo.14158810.

## Code availability

The Python package MEGA-FISH can be installed from PyPI (https://pypi.org/project/megafish/) using the “pip install megafish” command on Windows, Mac OS, and Linux systems. The GPU-accelerated version is available for Linux systems and Windows PCs with WSL2. Detailed instructions and API references can be found on ReadTheDocs (https://megafish.readthedocs.io/en/latest/index.html). The source code is hosted on GitHub at https://github.com/yumaitou/megafish and is distributed under the BSD-3-Clause License.

## Acknowledgements

We thank all members of the Ohkawa, Ochiai, and Nagasaki laboratories for helpful discussions and technical support. The infrastructure of the Omics Science Center Secure Information Analysis System, Medical Institute of Bioregulation, Kyushu University, provided part of the computational resources. We thank Margaret Biswas, PhD, from Edanz for editing a draft of this manuscript. This work was supported partially by JST FOREST JPMJFR2251 to K.T., JST CREST JPMJCR23N3 to H.O., JST NBDC JPMJDN2302 to M.N., Medical Research Center Initiative for High Depth Omics to Y.O., AMED BINDS JP22ama121017j0001 to Y.O., AMED CREST JP24gm1710012 to K.T., AMED JP23fk0210138 to K.T. and M.N., AMED JP21wm0425009, JP22fk0210111, JP22tm0424222, JP23ek0210194, JP23ek0109675, JP23ek0109672 to M.N., JSPS KAKENHI JP22H03538 to K.T., JP22H02609, JP22H04694, JP23H04286 and JP24H02326 to H.O., JP18H05527 to Y.I. and Y.O., JP23H00372, JP22H04676, JP22K19275 and JP24H02323 to Y.O.

## Author Contributions

Y.O. and Y.I. conceived the project. Y.O., Y.I., K.T., H.O. and M.N. designed the framework. Y.I. wrote the programs and analyzed the data. K.T., H.O. and M.N. provided materials. Y.O. and Y.I. wrote the manuscript with contributions from all authors.

## Declaration of competing interest

All authors declare no competing interests.

## Notes

https://github.com/yumaitou/megafish

### Competing Interest Statement

The authors have declared no competing interest.

