## Supplementary Information for "MEGA-FISH: multi-omics extensible GPU-accelerated FISH processing framework for huge-scale spatial omics"

| | MEGA-FISH | Big-FISH | RS-FISH | Starfish | pysmFISH* |
|---|---|---|---|---|---|
| **Calculation property** | | | | | |
| Processing image unit | Chunked whole image | Field of view | Field of view | Field of view | Mixed |
| Storing intermediate image | Zarr file | In memory | In memory | In memory (Can be output) | HDF5 file |
| Native multi-CPU core computation | Yes | No | No (Spark) | No (Batch job) | Yes |
| GPU computation | Yes | No | No | No | No |
| **Analysis methods** | | | | | |
| Image registration | Yes | No | No | Yes | Yes |
| Stitching | Yes | No | No | No | Yes |
| Segmentation | Yes | Yes | No | Yes | Yes |
| Importing cell labels | Yes | Yes | Binary mask | Yes | No |
| Spot decoding | Yes | No | No | Yes | No |
| SeqIS analysis | Yes | No | No | No | No |

**Supplementary Table 1** | Image processing frameworks for spatial transcriptomics. SeqIS, sequential immunostaining. *This framework was used in the EEL FISH analysis, Borm, L. E. *et al. Nat. Biotechnol.* **41**, 222–231 (2023).

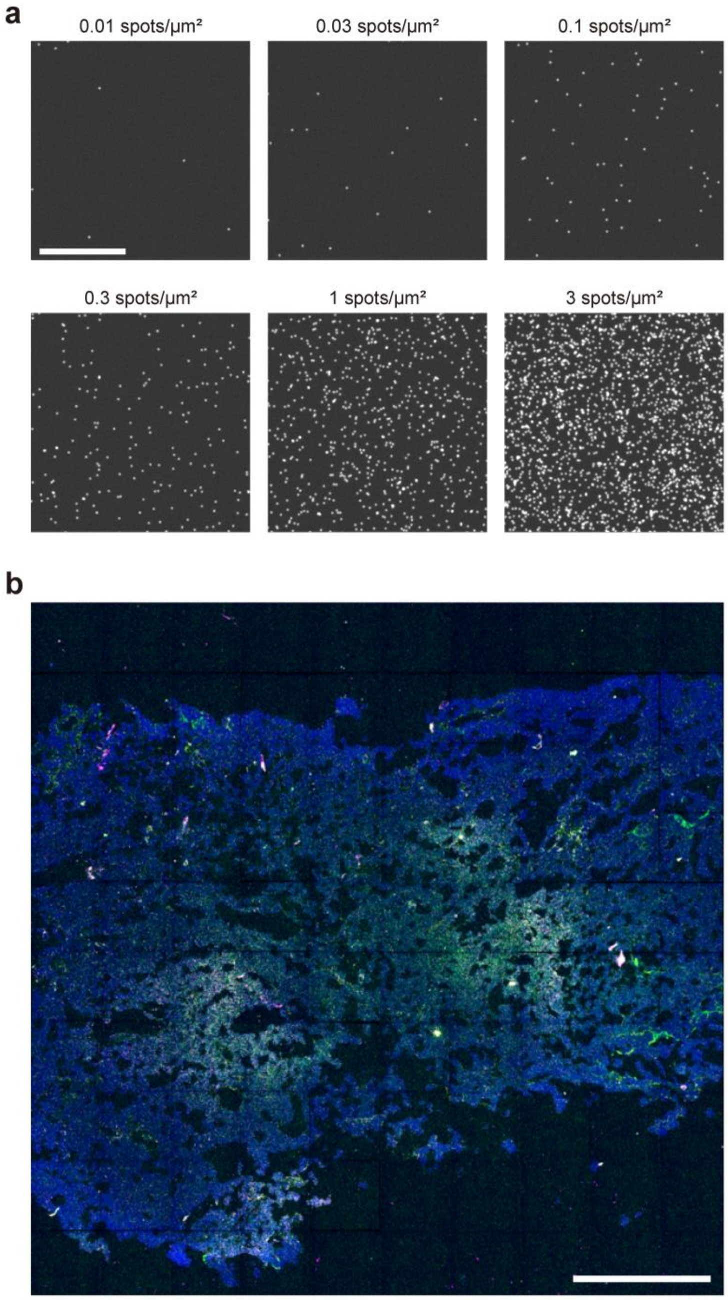


**Supplementary Figure 1** | Fluorescence In Situ Hybridization (FISH) images of simulation and tissue samples.

(a) Simulated RNA-FISH spots of various densities. Three-dimensional confocal point spread function images are displayed as maximum intensity projection along the z axis. Scale bar, 10 µm. (b) Representative RNA-seqFISH image of human carcinosarcoma tissue. The image shows a stitched view of the entire tissue section with Hoechst-stained nuclei (blue) and probes targeting *DHX36* (green), *HNRNPUL1* (magenta), and *ACIN1* (yellow). Based on Tomimatsu, K *et al.*, *Nat. Commun.* **15**, 3657 (2024). Scale bar, 400 µm.
